# Finding the balance between model complexity and performance: Using ventral striatal oscillations to classify feeding behavior in rats

**DOI:** 10.1101/241919

**Authors:** Lucas L. Dwiel, Jibran Y. Khokhar, Michael A. Connerney, Alan I. Green, Wilder T. Doucette

## Abstract

The ventral striatum (VS) is a central node within a distributed network that controls appetitive behavior, and neuromodulation of the VS has demonstrated therapeutic potential for appetitive disorders. Local field potential (LFP) oscillations recorded from deep brain stimulation electrodes within the VS are a pragmatic source of neural systems-level information about appetitive behavior that could be used in responsive neuromodulation systems. Here, we recorded LFPs from the bilateral nucleus accumbens core and shell (subregions of the VS) during limited access to palatable food across varying conditions of hunger and food palatability in male rats. We used standard statistical methods (logistic regression) as well as the machine learning algorithm lasso to predict aspects of feeding behavior using VS LFPs. These models were able to predict the amount of food eaten, the increase in consumption following food deprivation, and the type of food eaten. Further, we were able to predict whether the initiation of feeding was imminent up to 42.5 seconds before feeding began and classify current behavior as either feeding or not-feeding. In classifying this behavior, we found an optimal balance between model complexity and performance with models using 3 LFP features primarily from the alpha and high gamma frequencies. As shown here, unbiased methods can identify systems-level neural activity linked to symptoms of mental illness with potential application to the development and personalization of novel treatments.

**Author Summary:** As neuropsychiatry begins to leverage the power of computational methods to understand disease states and to develop better therapies, it is vital that we acknowledge the trade-offs between model complexity and performance. We show that computational methods can elucidate a neural signature of feeding behavior and show how these methods could be used to discover neural patterns related to other behaviors and used as therapeutic targets. Further, our results helps to contextualize both the limitations and potential of applying computational methods to neuropsychiatry by showing how changing the data being used to train predictive models (e.g., population vs. individual data) can have a large impact on how model performance generalizes across time, internal states and individuals.

## Introduction

The VS is a central node in the brain circuits influencing goal-directed and habitual behaviors with subregions like the nucleus accumbens (NAc) core and shell that mediate information between cognitive control regions of the prefrontal cortex and regions of learning and memory in the amygdala, hippocampus, and the ventral tegmental area [1-3]. As regions of convergence within the brain reward circuit, the NAc core and shell have been targeted by neuromodulation-based treatments for appetitive disorders including substance use disorders, eating disorders, and obesity [4-6]. Targeting the VS for eating disorders and obesity is supported by the known connectivity of the NAc with energy homeostasis circuits within the hypothalamus and brain stem [7,8] and the role of an-/orexigenic states modulating VS activity through central and peripheral ligands [9-11]. In both preclinical and clinical investigations DBS targeting these subregions has altered appetitive behavior [12-15], but variable treatment outcomes combined with the known risks of DBS have limited more widespread use. In other neuropsychiatric conditions, DBS outcomes have been improved and side-effects limited by using relevant neural activity to trigger stimulation, closed-loop DBS [16-18], or modify stimulation parameters, adaptive DBS [19]. However, a source of neural activity that contains information relevant to appetitive behaviors is needed to apply these advanced DBS approaches to appetitive disorders.

Local field potential (LFP) oscillations recorded from DBS electrodes are a promising source of neural systems-level information and VS oscillations could contain information about appetitive behavior. Although there has been evidence that high frequency components of NAc LFPs could be generated within the piriform cortex [20] the entrainment of single-unit firing to LFPs in the NAc suggests that VS LFPs are, at a minimum, a reflection of local striatal processing [21-23]. Therefore, it is likely that LFPs recorded from the NAc contain information pertaining to reward related behaviors.

Although a wealth of information likely exists within VS LFPs about appetitive behaviors like binge eating, it is important to determine how much information is needed to decode these behaviors and to determine what the trade-offs are between model performance and model simplicity. The model used by Wu et al. [18] took advantage of changes in delta power during the beginning of feeding in mice to trigger closed-loop DBS and reduced binge size. This is a good example of a low complexity population-based model (1 LFP feature trained from 6 animals) maximizing simplicity at the expense of performance (69% sensitivity and 64% specificity). Our work examined how altering model complexity changes performance in classifying feeding behavior in a rat model of binge eating. Unbiased statistical and machine learning approaches were used to predict imminent binge eating epochs and classify concurrent behavior using LFP recordings from bilateral NAc core and shell. Further, we used a machine learning approach to determine if VS oscillations recorded during binge eating sessions contained information that could predict the quantity of food consumed, the increase in binge size following food deprivation, or the type of food consumed (low vs. high palatability). Primarily, this work characterizes how different sets of training data (complex vs. simple; individualized vs. population-based) can affect the performance of models across individuals, time, and internal conditions. Although shown here in a rat model of binge eating, we expect these relationships to generalize to other appetitive behaviors and neuropsychiatric disorders, thus providing a framework for the development of more effective closed-loop algorithms and adaptive neuromodulation-based treatments.

## Materials and Methods

Male Sprague-Dawley rats (*n* = 12) were purchased from Charles River (Shrewsbury, MA) at 60 days of age were individually housed using a reverse 12-hour light/dark schedule with house chow and water available *ad libitum*. All experiments were carried out in accordance with the NIH Guide for the Care and Use of Laboratory Animals (NIH Publications no. 80-23) revised in 1996 and approved by the Institutional Animal Care and Use Committee at Dartmouth College.

### Experimental design

All animals were implanted with electrode arrays and were conditioned to binge eat using limited access to palatable food (1 month). Following completion of a DBS intervention published separately [24] and a 2 week washout period, rats re-established a baseline of binge-like feeding. Once a stable baseline was established, baseline binge sessions (Base) were recorded (LFP and video). Binge sessions were also recorded following 24- and 48-hour food deprivation (Dep 24 and Dep48) with intervening periods (at least 24 hours) of *ad libitum* access to house chow. Animals were also deprived of food for 24 hours before recording a 2-hour house chow session (Chow) to ensure that the animals consumed enough food to provide a comparable amount of LFP data during feeding to the three sessions with access to the high-fat/high-sugar diet (See General Methods: Surgery and Binge Eating).

These paired LFP and video recordings were used to address two types of questions: 1) **Whole session** questions—can VS activity averaged over an entire session be used to predict behavioral outcomes from that session (e.g., predict calories consumed)?; and 2) **Within session** questions—can VS activity be used to classify behavior as it occurs or to predict if it is about to occur? These two types of questions were assessed using tailored analysis approaches that have different sample size considerations and will be described in the corresponding subsections of Statistical Analysis.

### General methods

#### Surgery and binge eating

Following habituation to the animal facility, rats were anesthetized with isoflourane/oxygen (5% isoflourane for induction and 2% for maintenance) and underwent stereotaxic implantation with a custom electrode array targeting both the NAc core and shell bilaterally: 1.6 mm anterior; ±1 and 2.5 mm lateral; and 7.6 mm ventral to bregma. Following recovery from surgery (∼1 week), rats began a well described [25-27] schedule of limited access to a palatable high-fat/high-sugar diet (sweet-fat food), which contained 4.6 kcal/g and 19% protein, 36.2% carbohydrates, and 44.8% fat by calories (Teklad Diets 06415, South Easton, MA). The sweet-fat food was provided to the rats in addition to house chow and water within stimulation chambers for 2-hour sessions with 4-5 sessions irregularly spaced per week.

#### Behavioral measures

The food was weighed before and after all recording sessions to calculate the amount of food consumed during the session; weight was converted to kilocalories to allow direct comparison between food types (sweet-fat food vs. house chow). Session videos were manually scored for feeding, approach (movement towards food), and rest intervals. Timestamps were normalized by session length in order to compare feeding dynamics across sessions of unequal length (see Supplemental Methods: Behavioral measures).

#### Local field potential recording and processing

Rats were tethered in an 18”x12”x24” chamber through a commutator to a Plexon data acquisition system while time-synchronized video was recorded (Plexon, Plano, TX) for offline analysis. All LFP signal processing was done using custom code written from Matlab R2017a as previously reported [24] (see Supplemental Methods: Signal processing).

#### Verification of electrode placement

Rats were euthanized at the end of the experiment using CO_2_. Brains were removed, sectioned with a cryostat, mounted on slides, and stained with thionine for histologic verification of electrode placement [15]. No animals required exclusion based on electrode location.

### Statistical analysis - Whole session

To determine how much of the individual heterogeneity in feeding behavior could be predicted from VS activity, the values of each LFP feature were averaged across all bins according to behavioral scoring (i.e., feeding vs. resting) from a given session. To compare across animals and to account for day-to-day variation in signal fluctuations the LFP features were normalized by subtracting the average feeding value from the average rest value (Figure 2A). These rest-normalized LFP features were then used to predict behavioral variables (baseline food intake and food type). To predict the change in consumption with food deprivation—from Base to Dep24 or Dep48 session—the difference between the rest-normalized values of the two sessions were used as predictor variables (e.g., Base[feeding-rest] - Dep24[feeding-rest]). Although there were only 12 animals, by using up to two recordings per animal sample sizes were able to be increased (*n* = 24 for both baseline food intake and change in consumption with food deprivation; *n* = 21 for food type, one animal didn’t have any house chow sessions and another only had one); however, lasso is specifically for regressions in which p >> n and thus these samples size only limited the number of folds used in cross-validation.

The Matlab package *Glmnet* [28] was used to implement lasso with 100 iterations of 5-fold cross-validation. Since there were insufficient samples to construct a naïve test-set, model performance was estimated from the distributions of performances during cross-validation. For predicting baseline food intake and the change in consumption from baseline (continuous outcomes) performance was estimated as mean absolute error (MAE) and for predicting food type (binary outcome) performance was estimated as accuracy (%), both performance measures were reported with 95% confidence intervals. The performance distributions from cross-validation were then compared to distributions representing by-chance performance created by shuffling the assignment of predictor and outcome variables using Monte Carlo sampling. The actual and permuted distributions were compared using the Mann-Whitney *U* test and the *U* test statistic was converted into a Cohen’s *d* (Supplemental Methods: Effect size).

### Statistical analysis - Within session

Each 5-second bin and its corresponding LFP features belonged to only one behavioral category: feeding, pre-feeding, or not-feeding. Pre- and not-feeding were not manually scored; rather bins were assigned to those categories relative to the scored timestamps. Pre-feeding bins were defined as those occurring 45 seconds prior to feeding initiation and were discarded if they overlapped with a previous feeding epoch. All bins that were neither feeding nor pre-feeding were categorized as ‘not-feeding’.

#### Training pre-feeding vs. not-feeding models

Pre-feeding vs. not-feeding models were first trained and tested using pre-feeding bins that immediately preceded the start of feeding to determine if brain activity immediately before feeding was differentiable from brain activity during other behaviors. These models were then also tested on pre-feeding bins centered up to 45 seconds before feeding (Figure 3A). To account for the variability introduced through random sub-sampling and imputation of the bins used, all of these steps were repeated 20 times and the performance reported as averages with 95% confidence intervals (see Supplemental Methods: Pre-feeding vs. not-feeding).

#### Training feeding vs. not-feeding models

Logistic regressions, fit using the Matlab function *fitglm()*, were used to train models to classify bins as feeding or not-feeding (see Supplemental Methods: Feeding vs. not-feeding). The process of imputing and splitting the data into test and training sets was repeated 20 times for each condition (Base, Dep24, Dep48, and Chow).

#### Evaluating model performances

Model performances were measured using the naïve test-sets to calculate the classification probabilities of the new data with the function *predict()*, constructing a receiver operator characteristic (ROC) curve and calculating the area under the ROC curve (AUC) with the function *perfcurve()*. Performance was then averaged over the 20 iterations and reported with 95% confidence intervals. As before, Monte Carlo sampling used to shuffle the assignment of predictor and outcome variables in order to assess by-chance accuracy and the effect size between the distributions of actual and permuted performances was calculated by converting the *U* test statistic into a Cohen’s *d* (see Supplemental Methods: Effect size).

#### Relating model complexity to performance

The complexity of models used to differentiate brain activity during feeding from brain activity during not-feeding was adjusted across three domains: 1. the number of individual animals included; 2. the number of LFP features included; and 3. the number of conditions represented (Base, Dep24, Dep48, and Chow). Two-sample t-tests were used to compare the performance between models with different manipulations of the following domains followed by the Bonferroni correction.

##### 1. Individuals

To determine how many individual animals were required to create a population-based model that could generalize across animals, all permutations from 1/12 to 11/12 animals were used to train models. Population models (>1 animal used to train the model) were tested on the data of left-out animal(s) (see Supplemental Methods: Complexity vs. performance).

##### 2. LFP Features

The performance of the logistic model built from all LFP features was considered the “gold standard” for determining how many features were required to create a model that was stable across individuals and conditions. The lasso algorithm reduced the number of features used to ∼40. To test the lower bounds of simplicity, the performance of logistic models built from all possible permutations of up to three features: 58 monads, 1,653 dyads, and 30,856 triads. “Top-tier” models were those with performances not significantly different from the best performing model within each group as assessed using two-sample t-tests followed by the Bonferroni correction for multiple comparisons (i.e., top-tier monad models were monad models that did not perform significantly worse than the best monad model).

##### 3. Conditions

Models were trained using data from either just baseline or all four conditions, but in both cases more bins were used than needed to achieve stable model performance as determined by learning curves (Figure S2; see Supplemental Methods: Complexity vs. performance).

#### Visualizing LFP feature changes around feeding epochs

Features of interest (from monad, dyad, and triad models) were extracted from all recordings around feeding epochs from 62.5 seconds before feeding to 32.5 seconds into feeding, and from 12.5 seconds before feeding ended to 52.5 seconds after feeding ended (feeding epochs were only used if they lasted at least 45 seconds to avoid using data twice). For times outside of feeding, only data that did not overlap with previous or subsequent feeding epochs were used. The data from all baseline recordings were used to calculate the average LFP feature value in each 5 second bin around feeding epochs and were plotted through time with one standard deviation.

## Results

### Whole session results

#### Characterization of feeding dynamics across session types

There was a bimodal pattern of feeding behavior during the baseline conditions that was not maintained across the food deprived conditions (Figure 1A) in which more animals were feeding during the beginning of the recording. The temporal dynamics of feeding in animals given house chow after 24 hours of food deprivation was more similar to the deprived animals given palatable food rather than baseline; however, during the Chow condition the animals ate significantly fewer calories than both Dep24 and Dep48 conditions (*t*(19) = 3.73, p = .008, two-sample t-test; and *t*(16) = 5.82, p = .0002, two-sample t-test; Figure 1B). During the sweet-fat food sessions, 48 hours of food deprivation were required to significantly increase the number of calories consumed compared to baseline (*t*(19) = −4.25, p = .003, two-sample t-test; Figure 1B).

**Fig 1.**
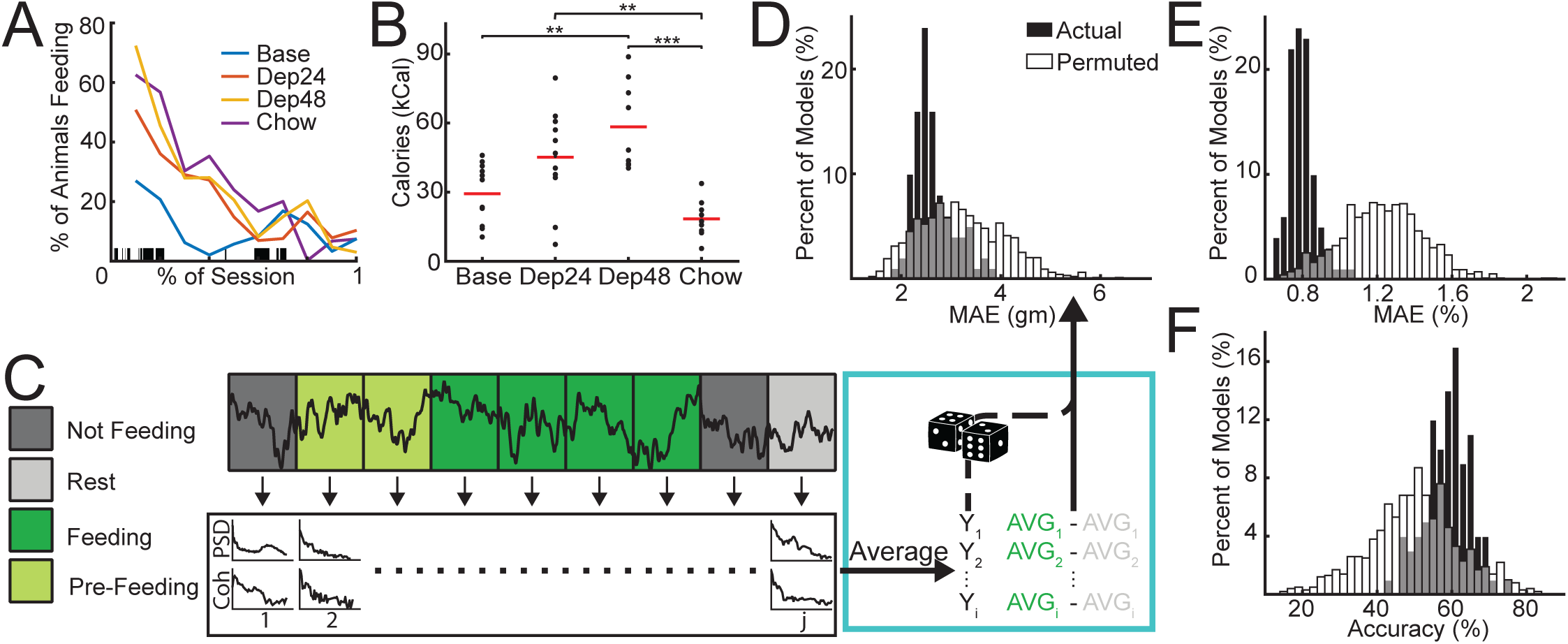
Predicting aspects of feeding behavior. **A.** Temporal dynamics of feeding behavior characterized on a population level across the four groups: baseline binge-like feeding of palatable chow (Base); binge-like feeding of palatable chow after 24-hours of food deprivation (Dep24) and 48-hours of food deprivation (Dep48); and limited access to house chow after 24-hours of food deprivation (Chow). Session lengths were normalized and the percent of animals feeding was smoothed for visualization. **B.** Kilocalories (kCal) consumed across conditions. More calories were consumed in Dep48 (*n* = 8) than Base (*n* = 12), and in both Dep24 (*n* = 12) and Dep48 than Chow (*n* = 8). Red bar indicates group average. * p < 0.05, ** p < 0.01 The change in kCal consumed were also used to calculate change in voracity (Figure 1-1A) which was then used to identify features potentially contaminated by chewing artifact (Figure 1-1B). **C.** Overview of analysis flow. Recordings were broken into non-overlapping 5 second bins and were assigned to one of 4 categories: not-feeding, rest, feeding, and pre-feeding. Power and coherence were calculated from each bin and averaged together according to category. Average brain activity of each animal during feeding was then normalized by activity during rest and used as predictor variables with behavioral metrics from that animal used as outcome variables. Outcome variables were then permuted to create distributions of ‘by-chance’ performances. **D-E.** Performance of models in predicting behavioral metrics; black distributions are performances of models using actual outcome variable assignment and white distributions are performances of models using permuted outcome variables. Distribution statistics reported as mean±95% confidence interval and effect size between Actual and Permuted given as Cohen’s *d*. **D.** Mean absolute error (MAE) of predicting baseline food consumption in grams (gm). Actual = 2.6±0.08 gm; Permuted = 3.2±0.05 gm; *d* = 0.43. **E.** MAE of predicting percent change (%) in food consumption in Dep24 and Dep48 compared to Base (%): Actual = 0.8±0.01%; Permuted = 1.2±0.01%; *d* = 1.1. **F.** Percent accuracy (%) in predicting if house chow or palatable food was being consumed: Actual: 59±1%; Permuted: 50±1%; *d* = 0.52.

**Fig 2.**
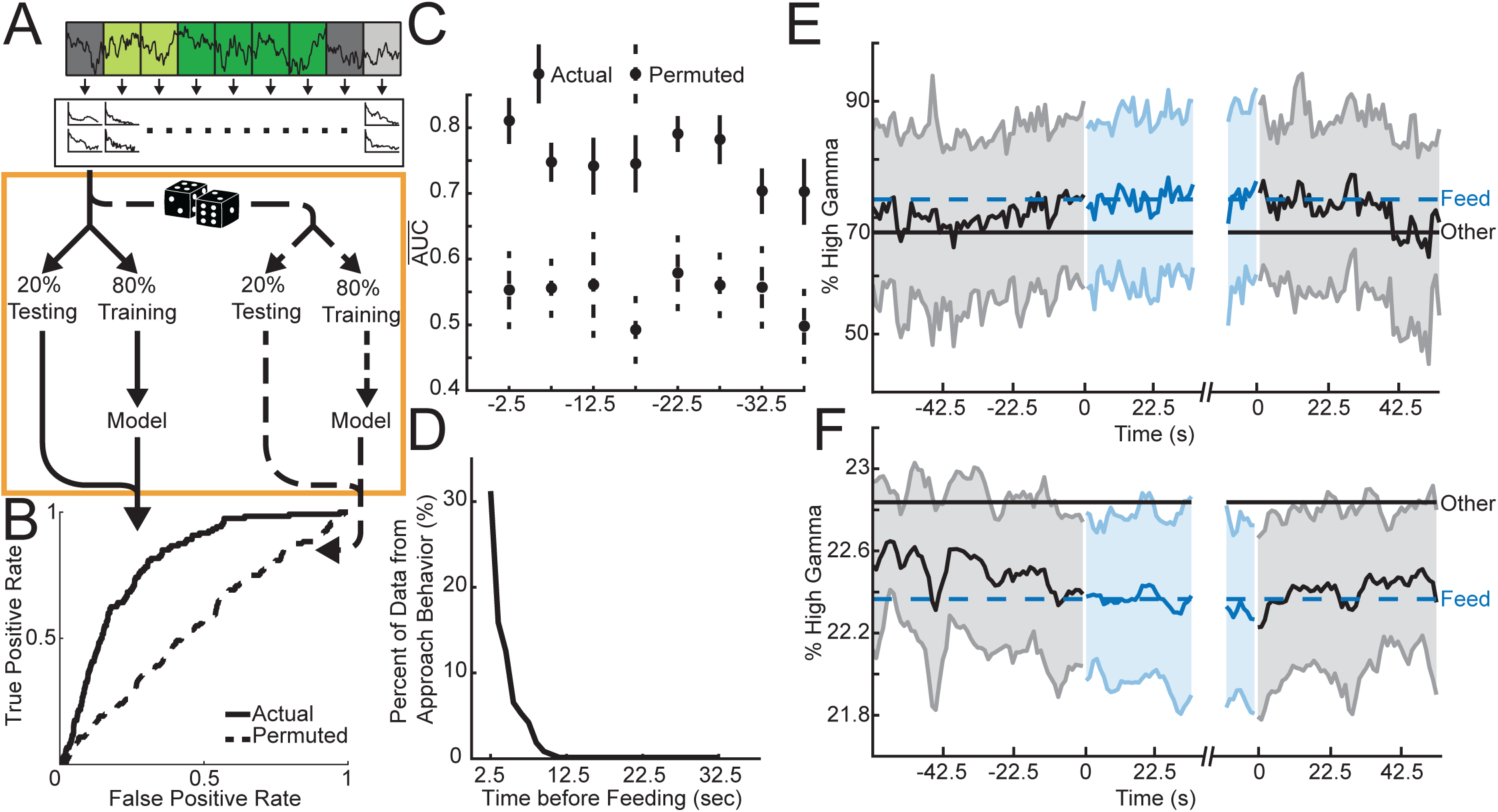
Classifying imminent feeding from other behavior with VS oscillations. **A.** Overview of analysis flow. Data was binned and power and coherence extracted as in Figure 1. Orange box: Eighty percent of all pre-feeding bins immediately before feeding bins and all other bins outside of feeding (not-feeding and rest) are used for model training and the other 20% are used for testing. Using the same data, the assignment of category to predictor variables is permuted and trained on 80% of the data and tested on the left out 20%. This analysis produces two receiver operator (ROC) curves representing the performance of models built from actual and permuted data. To build rare event detectors (feeding is only ∼18% of the data) the adaptive synthetic sampling approach for imbalanced learning was found to be the best option (Figure 2-1A-B). To be able to compare across models it was also necessary to determine the number of samples were required to achieve stable model performance (Figure 2-1A-D). **B.** Average performance of differentiating pre-feeding immediately before feeding begins and bins outside of feeding (not-feeding and rest); performance distribution statistics reported as AUC mean ± 95% confidence interval and the effect size between Actual and Permuted given as Cohen’s *d*: Actual = 0.81±0.03; Permuted = 0.55±0.06; *d* = 2.68. **C.** Average areas under the ROC curves 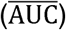 obtained for 20 iterations of model building using models built from the pre-feeding bins immediately preceding feeding and tested on all other pre-feeding bins grouped by amount of time before feeding up to bins centered 42.5 seconds before feeding. **D.** Percent of pre-feeding data which overlaps with approach behavior; pre-feeding bins centered up to 12.5 seconds before feeding have at most 30% overlap with approach behavior. **E-F.** Normalized representative LFP features plotted through time around feeding. Features were normalized by either total power or by average coherence of the channel(s) of interest and averaged across all bins and animals around the beginning and end of feeding epochs. Shading indicates ±1 standard deviation. Black traces represent averages before and after feeding and blue traces represent averages during feeding. Dashed blue and solid black lines indicate average feature activity for either feeding bins or all other bins including those outside of these plots. **E.** Core left core right high gamma coherence increases during feeding. **F.** Shell left high gamma power decreases during feeding. Although both **E** and **F** exhibit a mean shift, a decrease in feature variance around feeding was also found (Figure 2-2).

#### Decoding the amount of food consumed and food type from VS oscillations

Using the brain activity at rest recorded during baseline sessions (Figure 1C) models were built that outperformed permuted data in predicting the amount of food eaten at baseline (MAE = 2.6±0.08 gm; *d* = 0.43; Figure 1D). Next, changes in brain activity at rest from Base to Dep24 or to Dep48 were used to predict the changes in the amount of food consumed across the corresponding sessions (MAE = 0.8±0.01%; *d* = 1.1; Figure 1E). Last, brain activity at rest was used to predict the type of food being eaten following 24-hour food deprivation—house chow vs. sweet-fat food (accuracy = 59±1%; *d* = 0.55; Figure 1F).

### Within session results

#### Predicting imminent feeding

LFP features from the bins immediately preceding feeding epochs in the baseline session were used to build models (Figure 2A) that predicted whether initiation of feeding was imminent (AUC = 0.81±0.03; *d* = 2.68; Figure 2B). When these models were applied backwards through time, they were able to successfully differentiate bins as preceding feeding (i.e. pre-feeding bins centered up to 42.5 seconds before feeding began) from all other bins outside of feeding epochs (Figure 2C). These models were not merely detecting approach behavior since only one third of the pre-feeding bins had any overlap with approach behavior and none of the bins more than 12.5 seconds before feeding overlapped with approach while the model was still able to differentiate beyond 12.5 seconds (Figure 2D).

The identity of the LFP features that were most frequently used by lasso to predict imminent feeding were plotted over time to investigate the temporal link between changes in these LFP features and feeding behavior. Both a power (shell left high gamma; Figure 2E) and coherence feature (core left-core right high gamma; Figure 2F) tended to transition from the non-feeding average towards the feeding average during the pre-feeding interval (42.5 seconds before feeding). After feeding ceded, the LFP features began to transition back to their non-feeding level.

### Relating model complexity to performance using models classifying feeding vs. not-feeding

To explore model complexity, three domains of the data used for model building were manipulated (Figure 3A): number of individuals, number of LFP features, and number of conditions (Base, Dep24, Dep48, and Chow). By using the same naïve test-sets for assessing performance, the effects of different training data could be directly on performance could be compared directly.

**Fig 3.**
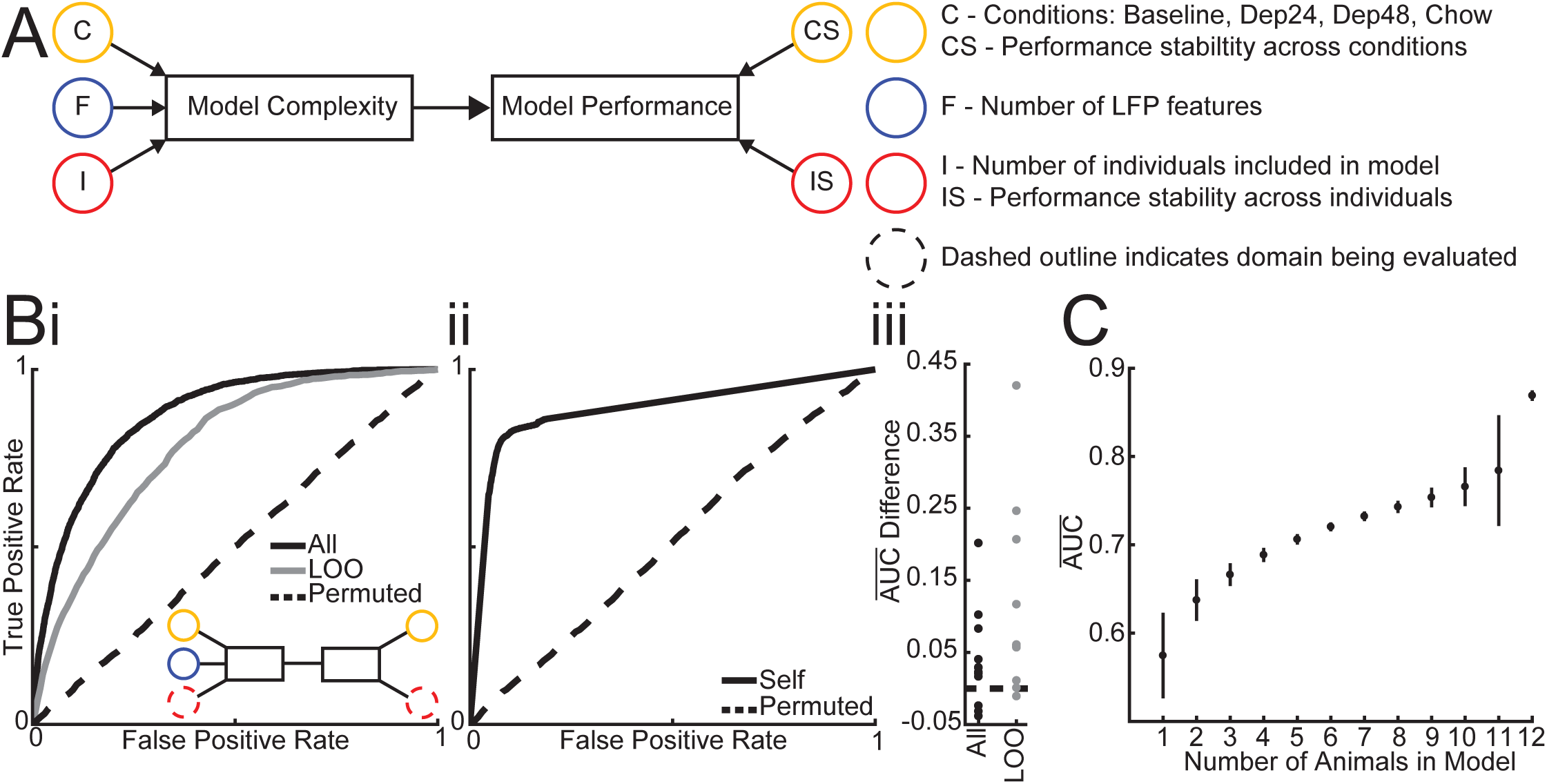
Number of individuals needed to build models. **A.** Schema of factors influencing model complexity and affected facets of model performance. Dashed outline indicates that a given factor is being manipulated in current analysis and that the performance is being measured accordingly. **B.** (Inset) The number of individuals and performance stability across individuals is assessed by comparing the average performance of population models (i) built using 80% of all animals’ data and tested on the remaining 20% (All = 0.87±0.01; *d* = 3.27), using a leave-one out (LOO = 0.78±0.06; *d* = 3.13) approach, and using permuted data in a leave-one out approach (Permuted = 0.51±0.02) to the average performance of individual models (ii) built from 80% of each animals’ data and testing on that animals’ left out 20% (Self = 0.89±0.06; *d* = 3.13) alongside permuted individual models (Permuted = 0.52±0.02). (iii) The subtraction of individual model performance on classifying the left out data of that individual from the performance of population models on the same data shows that individual models almost always outperform population models, using both All and LOO datasets. **C.** Average performance of models built from all possible combinations of including 1 to 12 individuals and tested on population data. Circle indicates mean and vertical bar indicates 95% confidence interval.

#### Manipulation of the number of animals in population-based models

Population-based models built from and tested on data from all animals was able distinguish brain activity during feeding from not-feeding better than would be expected by chance alone (AUC = 0.87±0.01; *d* = 3.27; Figure 3Bi). All models were built using 80% of each individual animal’s data from baseline sessions (Figure 2A) and the models were tested on the left-out 20%. However, it is conceivable that because the left-out 20% came from the same recordings as the 80% used for training, these models could be slightly overfit. To account for this a leave-one-out (LOO) approach was used by leaving an entire animal out of the model building to be used as the test-set. As expected, the performance of LOO models decreased significantly (t(258) = 2.96 p = .003, two-sample t-test), but they still outperformed the by-chance permuted models (AUC = 0.78±0.06; *d* = 3.13; Figure 3Bi). As the number of animals left out was increased (Figure 3Bi Inset) the performance continued to decline (Figure 3C).

#### Individualized vs. population-based models

The above data indicate that if enough individuals are used to train population-based models, then the model will generalize across individuals, outperforming chance. It is possible however that a model built from an individual will be able to perform better for that individual compared to a model built from a population. To test this individualized models were built using 80% of a single animal’s data and then tested on that animal’s left-out 20%. These models performed better than the permuted models (AUC = 0.89±0.06; *d* = 3.13; Figure 3Bii) and performed better than the population-based models tested on the same left-out data in 9/12 individuals when all animals were in a population model (Figure 3 Biii All) and 11/12 individuals when all but that individual was used to train the population-based model (Figure 3Biii LOO).

#### Manipulation of the number of LFP features at baseline

The next step was to determine how many LFP features are necessary to build successful and stable models by manipulating the number of features included in model building from all single feature models to all three feature models (Figure 4A Inset). When classifying baseline feeding vs. not-feeding, the best models built from a single feature performed better than chance and although increasing the number of features used to two or three increased performance, these simple models never achieved the performance of either the lasso (∼40 features) or full logistic (58 features) model (Figure 4A). Further, when these simpler models were applied to testing data from the other conditions (Dep24, Dep48, and Chow) the performance dropped with the more complex triadic models providing the best performance stability across conditions, approaching the stability of the full logistic models (Figure 4B). Last, the features used by the best dyad and triad models and those used by lasso were not all top performing features when used in monadic models, suggesting redundancy between LFP features in predicting feeding behavior (Figure S4).

**Fig 4.**
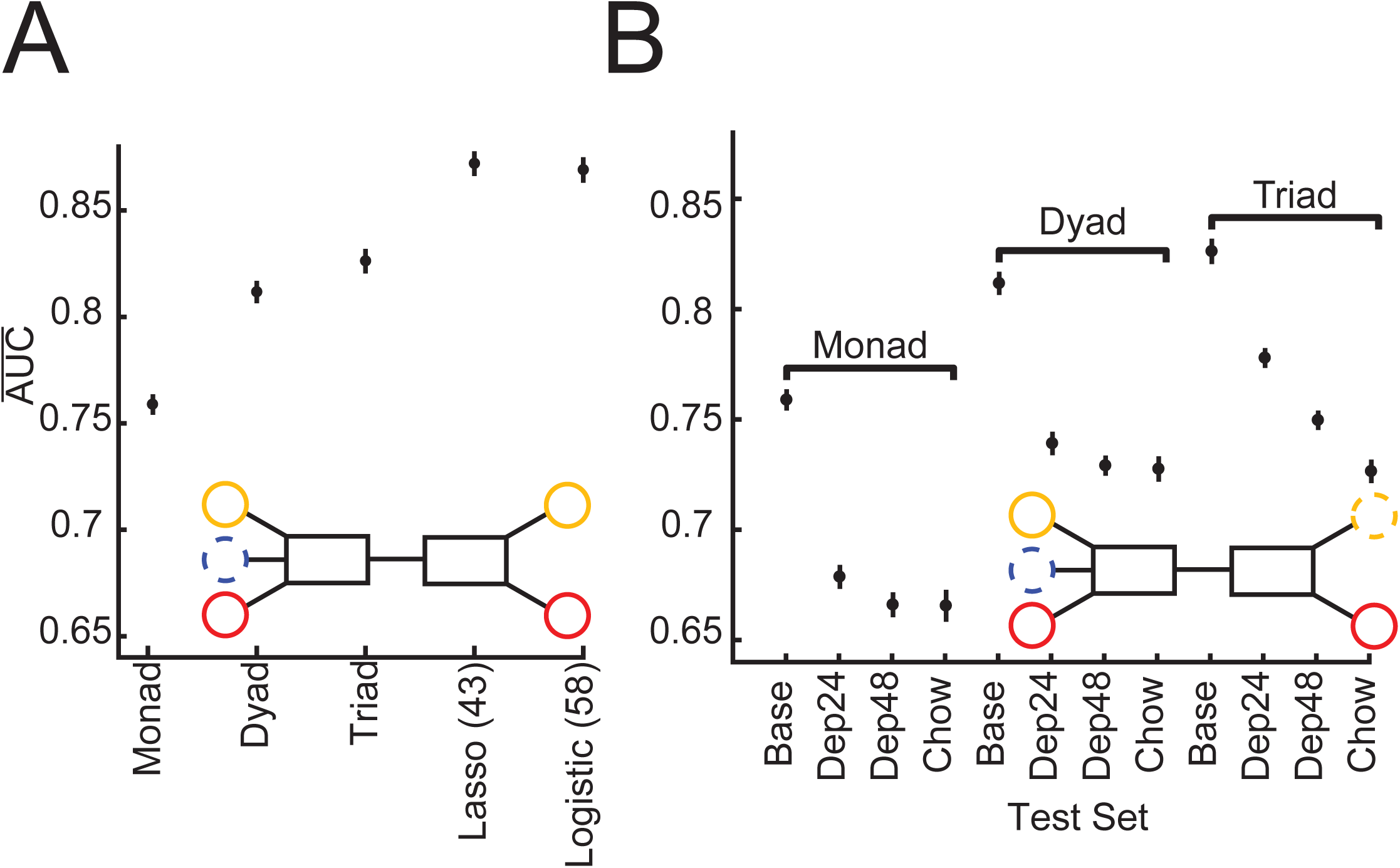
Number of LFP features needed to classify feeding. **A.** Average performance of models built and tested on population baseline data as the number of features used is manipulated (Inset): single features (Monad), two features (Dyad), and three features (Triad), using lasso (Lasso), and all 58 features (Logistic). The top dyad and triad were not composed of the top two and three monad models, suggestive of the redundancy of LFP features (Figure 4-1). Average performance of all population baseline Monads, Dyads, Triads, and full Logistic across conditions (Inset). Averages represented by circles and 95% confidence intervals by vertical bars.

#### Manipulation of conditions used to train and test models: impact on individual and population model performance

Given that individualized models tended to outperform population-based models when trained and tested in the baseline condition the performance of these models was also determined when data from the other three conditions were used as the test-set. The population-based models significantly outperformed the individual models (Dep24 *t*(26) = −4.03, p = .0035; Dep48 *t*(26) = −3.53, p = .012; and Chow *t*(26) = −3.63, p = .0097; Figure 5A). To determine if the model performances could be improved across the conditions, new individualized and population-based models were built using data from all four conditions (Figure 5A Inset). These individualized models significantly outperformed the population-based models (Base *t*(26) = 3.56, p = .012; Dep24 *t*(26) = 3.94, p = .0044; and Dep48 *t*(26) = 9.69, p = 3.3E-9; Chow *t*(26) = 3.52, p = .013; Figure 5B). In summary, the population-based models built from the baseline condition generalized better across the other conditions compared to the individualized models. However, if individualized models are trained on data from all conditions, then they can outperform the population-based models in classifying feeding vs. not-feeding across conditions.

**Fig 5.**
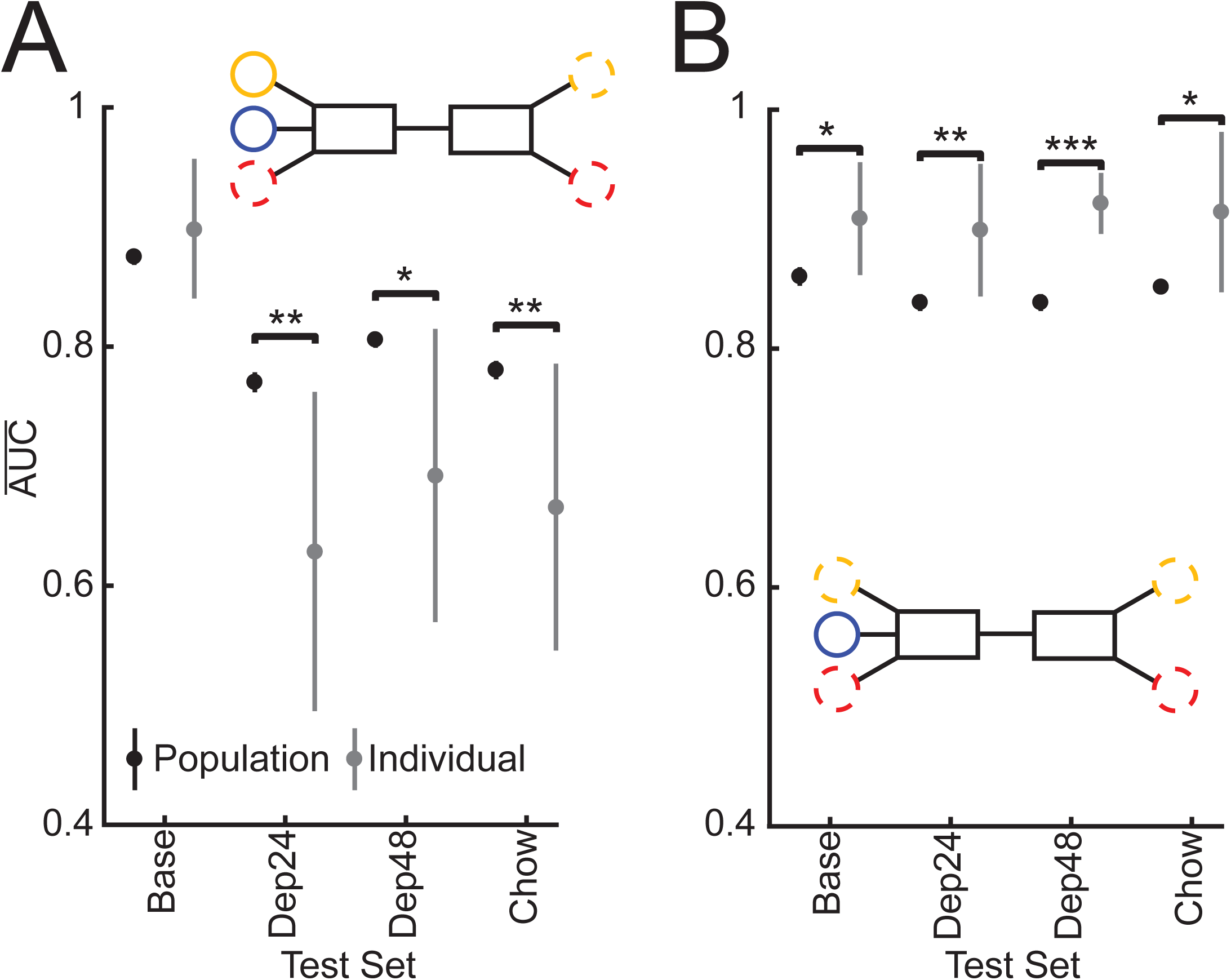
Performance of Population (black) and Individual (grey) models built from Baseline (**A**) and all condition (**B**) data and tested across states. **A.** Testing performance of population and individual models built from baseline data on all conditions (Inset). Population models built from Base do better than individual models when predicting behavior during any other condition than Base. **B.** Testing performance of population and individual models built from all conditions on all conditions (Inset). When models are built from all conditions, individual models do better than population models across conditions. * p < 0.05, ** p < 0.01, *** p < 0.001. Averages represented by circles and 95% confidence intervals by vertical bars.

#### Manipulation of the number of LFP features used to build models across conditions

Finally, the above findings were integrated by determining the minimal number of LFP features required in an individualized model built from all conditions to match the performance of the corresponding, “gold standard”, logistic model using all 58 LFP features (Figure 6 Inset). Testing on Base and Chow data revealed that only two features were required to attain a performance that was not significantly different from the gold standard logistic model (Base *t*(14) = 3.07, p = .13; Chow *t*(14) = 1.74, p = 1; Figure 6A and D). The Dep24 condition required 48 features used by lasso (Dep24 *t*(14) = 3.19, p = .10; Figure 6B) to perform at a comparable level to the full logistic. However, testing all possible combinations of LFP features from 4 to 48 was beyond the computational capacity of this study.

**Fig 6.**
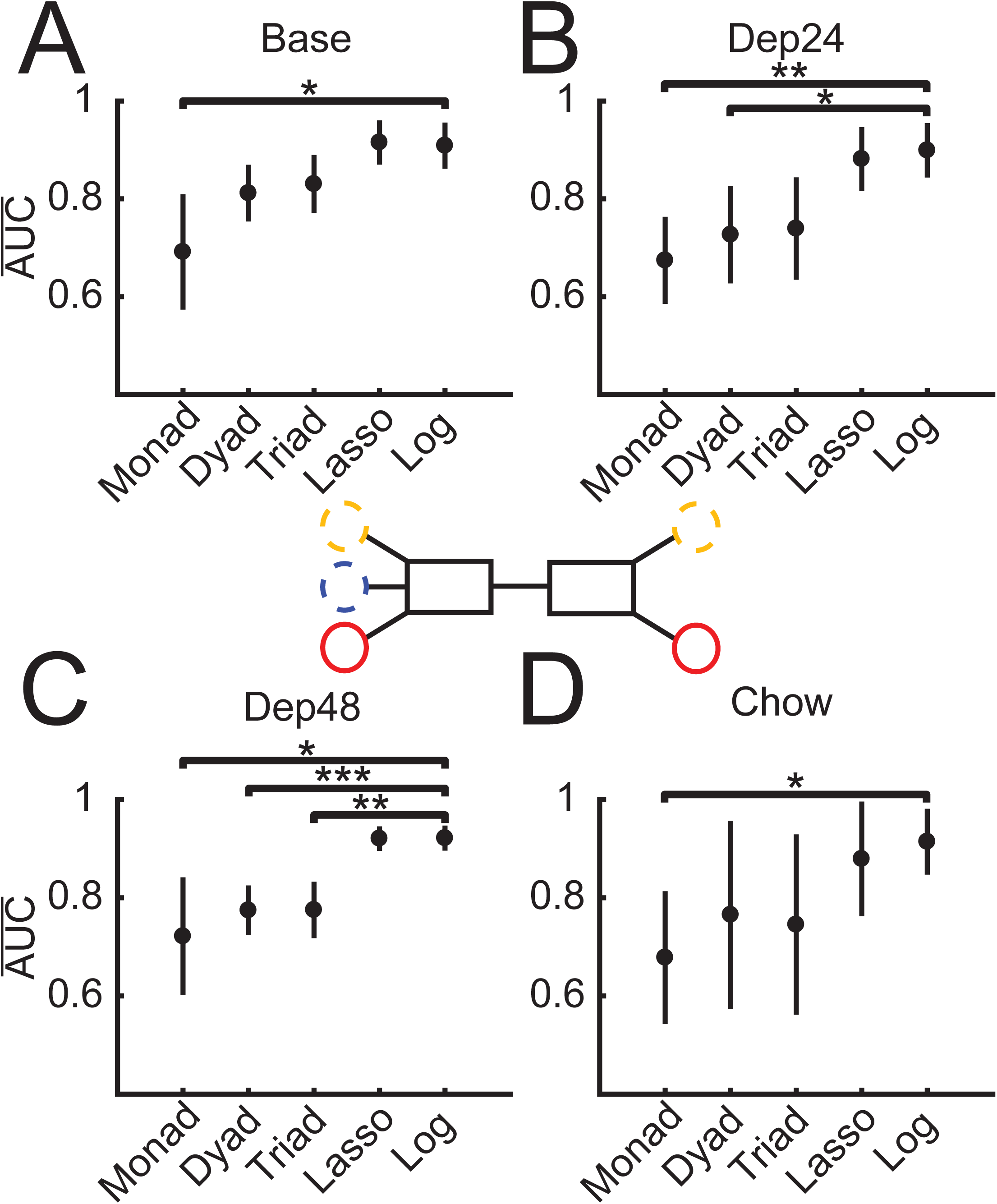
Manipulating all number of features and using all conditions to find simplest individualized models (Inset) and using the full 58 feature logistic regression (Log) as the performance to match. For predicting behavior at Base (**A**) and Chow (**D**) two features are sufficient to achieve maximal performance. For Dep24 (**B**) three features are sufficient and for Dep48 (**C**) Lasso is required. **A.** Only Monads have significantly lower performance than the full Log, *t*(14) = 4.04, p = .019. **B.** Monads, *t*(14) = 5.07, p = .0027, and Dyads, *t*(14) = 3.58, p = 0.049, p = have significantly lower performance than Log. **C.** Monads, *t*(14) = 3.85, p = .028, Dyads, *t*(14) = 6.12, p = .00042, and Triads, *t*(14) = 5.51, p = .0012, have lower performance than Log. **D.** Only Monads have significantly lower performance than Log, *t*(14) = 3.70, p = 0=.038. * p < .05, ** p < .01, and *** p < .001. Averages represented by circles and 95% confidence intervals by vertical bars.

When data from all conditions were used from training and testing it was found that many of the models performed equally well (Table S1). Despite the flexibility in the exact features used in these models, features within the alpha and high gamma ranges were the most common contributors to the top performing models (Table 1). For the simpler models—monads and dyads—most predictors were power features while in the more complex triads coherence features became equally represented (Table S1).

**Table 1.** Appearance of frequency ranges in top-tier models (models that are not significantly worse than the top performer). There are 2 Monads, 8 Dyads, and 32 Triads within the top-tier (Table 1-1). Percentages of dyads and triads do not add to 100% since each model can contain more than one frequency range.

|  | Monad | Dyad | Triad |
| --- | --- | --- | --- |
| Delta | — | 38% | 44% |
| Theta | — | — | — |
| Alpha | 50% | 88% | 84% |
| Beta | — | 13% | 13% |
| Low Gamma | — | — | 44% |
| High Gamma | 50% | 63% | 72% |

Table 1: Frequency contributions in top performing monad, dyad, and triad models.

## Discussion

Here we show that VS oscillations can be decoded and used to predict aspects of feeding behavior such as the amount and type of food consumed. Further, these oscillations can be used in real-time to predict imminent or current feeding in a rat model of binge eating, regardless of varying hunger levels and food palatability. By manipulating the complexity of the training data used and then assessing how these manipulations affect model performance, this work creates a theoretical framework to guide the implementation of predictive algorithms used in closed-loop and adaptive neuromodulation systems. Specifically, this work suggests that devices could be pre-loaded with a model built from a group of individuals using a handful of oscillatory features and have fair performance when applied to individuals outside of the training group. However, to achieve optimal performance in a specific individual, personalized models using neural data from that individual acquired over time would be needed.

### Predicting amount of food consumed and food type from VS oscillations

The binge eating model used in this study resulted in variation in food intake between individuals (as previously described, [29]) and also in the increased intake following food deprivation. That this variation could be partially predicted using LFP features from the VS is not surprising given the connectivity of these VS regions (e.g., nucleus accumbens) to the distributed feeding network [30,31]. Further, it is feasible that information about the type of food consumed (low vs. high palatability) could also be extracted from VS oscillations given that ‘liking’ of food [32,33] and nutritive/caloric value [34] are processed by the NAc.

### Population-based vs. individualized models

The presented data suggest that models classifying feeding behavior perform best when they are built and tested within a given individual and condition, otherwise performance drops when these individualized models are tested in other conditions—even if within the same animal. However, this drop can be mitigated in two ways: 1. use data from multiple conditions within an individual; or 2. use data from multiple individuals within a condition (or from multiple conditions). Following either of these strategies will increase the probability of selecting and properly weighting LFP features that generalize across animals and conditions. Clinically, it is important that models are able to perform well across a heterogeneous population as well as across the variable conditions through which individuals will transition over time. Therefore, when building models for use in algorithms that control closed-loop or adaptive neuromodulation systems it will be vital to consider these strategies.

### Machine learning and DBS

Recently, there has been increasing interest in applying machine learning and other computational methods to guide the selection and implementation of psychiatric treatments. This work can be categorized into three domains: 1. predicting treatment outcomes, 2. defining treatment parameters, and 3. actively optimizing treatment through time. Treatment outcomes have been successfully predicted using both structural and functional connectivity measures in Parkinsonian patients [35] as well as using LFPs in a rodent model of binge eating [24]. In a high throughput manner, machine learning has also been used to optimize combinations of DBS parameters and medications [36] and to determine which DBS parameters lead to desired changes in brain activity [37]. Last, there is a push towards the development of neuromodulation systems that use machine learning to monitor stimulation evoked dopamine signaling and adjusting stimulation parameters to optimize treatment [38] or to determine the time and brain state during which stimulation has the largest effect [39]. With the exception of Kumar et al.’s work, all of these implementations of machine learning used population-based datasets (i.e. data pooled across subjects). For some predictions—e.g., optimizing stimulation target—using individual datasets is impractical; however, when tuning stimulation parameters or trying to determine effective stimulation timing our work suggests individualized data will produce the best models, if those data were sampled from across time and, ideally, across varying conditions (e.g., stress/anxiety, mood, hunger, etc.).

### Number of predictors required for stable performance

This work highlights an important trade-off between the number of features used to build a model and that model’s performance across individuals and conditions. Our results suggest that single feature models are unlikely to perform with enough accuracy to be clinically useful. However, a relatively simple model (∼3 features) can approach the performance and stability of complex models (58 features); although these exact numbers may vary when predicting different behaviors/symptoms, this work suggests that simpler models can contain enough information pertaining to a given behavior/symptom with the benefit of limiting the computational and power requirements.

### A predictable pattern of brain activity precedes feeding initiation

As shown (Figure 2), VS oscillations can differentiate times when an animal is about to feed (≤45 seconds before feeding begins) from not-feeding. Interestingly, models built to classify feeding were also able to classify pre-feeding—even slightly outperforming pre-feeding models. This was because although many of the same LFP features were used in both models, the features are more clearly delineated (i.e., higher signal to noise ratio) between feeding and not-feeding, likely representing a “ramping” of LFP features as feeding begins.

When this hypothesis was examined (Figure 2E and F), a ramping phenomenon was observed in features that more successfully classified pre-feeding bins when trained on feeding vs. not-feeding data. For example, NAc shell left high gamma power begins to increase to its feeding level before feeding begins, stabilizes during feeding, then drops back to not-feeding levels as feeding ends. In other features there is decreased variance during feeding (Figure 2-2), mirroring a widespread cortical pattern seen in response to obtaining rewards [40], as a marker of stimulus perception/detection [41,42], and has been correlated with performance on visual discrimination [43,44] and motor preparation/initiation [45,46]. Visualizing several of these LFP features over time makes it clear that they develop over different timescales, likely underpinning the accuracy of predicting imminent feeding increasing closer to the start of feeding as more features have diverged from their not-feeding levels (Figure 2C).

### Confounds and noise

Given that motor preparation and initiation manifest as decreased neural variability it is possible that the models predicting imminent feeding were merely detecting the motor activity of approaching the food. However, by 12.5 seconds before feeding none of the data used for model building contained approach behavior (Figure 2D) and the models were able to classify pre-feeding brain activity out to 42 seconds before feeding without a corresponding drop in performance at 12.5 seconds (Figure 2C). When models were built using feeding data—with no approach behavior—performance was actually slightly better at classifying pre-feeding. Together these results suggest that although approach behavior may also exhibit a detectable change in LFPs, this change is not a primary source of information for our models.

There is also a concern that without electromyography of the masseters we are unable to completely remove the possibility of chewing noise influencing models built from feeding data. However, if the feeding vs. not-feeding models were chewing detectors, then the models should only be able to detect behaviors with chewing noise. Instead the feeding models were also able to classify pre-feeding activity which is devoid of chewing noise.

### VS oscillations as a source of information for closed-loop and adaptive neuromodulation

Appetitive disorders like binge eating disorder have been associated with dysregulation of the brain reward circuit which includes the VS. Understanding the nature of network pathology is critical to the development of treatments that can therapeutically alter network activity, ameliorating the problematic behavior/symptom. Neuromodulation-based interventions that use network activity to trigger stimulation and modify parameters have proven effective in the treatment of Parkinson’s disease [19]. DBS for epilepsy has incorporated electrodes in order to detect pre-seizure electrophysiologic activity and use stimulation to prevent seizure generation [16]. Future treatments of appetitive disorders could similarly utilize meaningful electrophysiology features in order to provide feedback to optimize treatment efficacy and limit the side effect profile. A recent study demonstrated the theoretical feasibility of using a closed-loop system to trigger stimulation and decrease binge size in a mouse model of binge eating [18]. As noted in the introduction, the model used had limited performance, likely due to only using a single LFP feature, but the success of the intervention highlights another point to consider when determining model complexity vs. model performance tradeoffs: what model performance is required to meaningfully impact the outcome of interest?

### Conclusions

Ventral striatal oscillations can predict feeding behavior during sessions of limited access to palatable food in rats. Similar unbiased computational methods will continue to identify new systems-level neural markers and produce models capable of predicting behaviors that are relevant to an array of neuropsychiatric conditions.

---

**Fig S1**. Features potentially contaminated by chewing noise **A.** Distribution of percent changes in voracity from baseline to food deprived conditions (Dep24 and Dep 48). **B.** Regression between percent change in shell left core left theta coherence and percent change in voracity from baseline to food deprived conditions; p < 0.01; R^2^ = 0.3.

**Fig S2**. Learning curves used to determine minimum number of trials required to make models whose performance stability are comparable. Circles indicate average AUC and vertical bars represent 95% confidence intervals. Grey boxes indicate at which trial number performance becomes insignificantly different from performance of models with most data. **A.** Testing population models using ADSYN imputation and between 6 and 500 trials per animal; 150 per animal (1800 total) are needed to achieve performance of models using all 500 per animal (6000 total). **B.** Testing population models using weighted outcome variables and between 6 and 500 trials per animal; 350 trials per animal (4200 total) would be needed to match performance of full models. **C.** Testing individual models (grey lines) using between 30 and 500 trials; the most any individual needed was 350 trials to match the performance of models made with all 500 trials. **D.** Testing pre-feeding models using between 4 and 350 trials each; 13 trials per animal were needed to match the performance of 350 trials per animal.

**Fig S3**. Monad feature performance with features used for top performing Dyad and Triad indicated and features with 100% survival in the top performing Lasso models highlighted in red. Average AUC indicated by a circle and 95% confidence interval by a vertical bar.

**Fig S4**. Shell right low gamma power exhibits a marked decrease in variation during feeding. Power was normalized by total power of shell right and averaged across all trials and animals around the beginning and end of feeding epochs. Shading indicates ±1 standard deviation. Black traces represent averages before and after feeding and blue traces represent averages during feeding. Dashed blue and solid black lines indicate average feature activity for either feeding trials or all other trials outside of these plots.

**Table S1.** Top-tier monads, dyads, and triads with features and performances (area under the curve, AUC). Locations: Sell left (SL), shell right (SR), core left (CL), core right (CR). Frequencies: delta (d), theta (t), alpha (a), beta (b), low gamma (lg), and high gamma (hg). Power features have one location and frequency (e.g. shell right alpha power is SRa) and coherence have two locations and one frequency (e.g. high gamma coherence between shell left and core right is SLCRhg). Feature order (Feature 1 vs. Feature 2) is not important.

## References

1. Berridge KC, Ho CY, Richard JM, DiFeliceantonio AG (2010) The tempted brain eats: pleasure and desire circuits in obesity and eating disorders. Brain Res 1350: 43–64.

2. Ferrario CR, Labouebe G, Liu S, Nieh EH, Routh VH, et al. (2016) Homeostasis Meets Motivation in the Battle to Control Food Intake. J Neurosci 36: 11469–11481.

3. Volkow ND, Wang GJ, Fowler JS, Tomasi D, Baler R (2012) Food and drug reward: overlapping circuits in human obesity and addiction. Curr Top Behav Neurosci 11: 1–24.

4. Muller UJ, Voges J, Steiner J, Galazky I, Heinze HJ, et al. (2013) Deep brain stimulation of the nucleus accumbens for the treatment of addiction. Ann N Y Acad Sci 1282: 119–128.

5. Pierce RC, Vassoler FM (2013) Deep brain stimulation for the treatment of addiction: basic and clinical studies and potential mechanisms of action. Psychopharmacology (Berl) 229: 487–491.

6. Kumar R, Simpson CV, Froelich CA, Baughman BC, Gienapp AJ, et al. (2015) Obesity and deep brain stimulation: an overview. Ann Neurosci 22: 181–188.

7. Carus-Cadavieco M, Gorbati M, Ye L, Bender F, van der Veldt S, et al. (2017) Gamma oscillations organize top-down signalling to hypothalamus and enable food seeking. Nature 542: 232–236.

8. Leigh SJ, Morris MJ (2016) The role of reward circuitry and food addiction in the obesity epidemic: An update. Biol Psychol.

9. Berendse HW, Groenewegen HJ, Lohman AH (1992) Compartmental distribution of ventral striatal neurons projecting to the mesencephalon in the rat. J Neurosci 12: 2079–2103.

10. Zahm DS, Brog JS (1992) On the significance of subterritories in the “accumbens” part of the rat ventral striatum. Neuroscience 50: 751–767.

11. Groenewegen HJ, Wright CI, Beijer AV (1996) The nucleus accumbens: gateway for limbic structures to reach the motor system? Prog Brain Res 107: 485–511.

12. Vassoler FM, Schmidt HD, Gerard ME, Famous KR, Ciraulo DA, et al. (2008) Deep brain stimulation of the nucleus accumbens shell attenuates cocaine priming-induced reinstatement of drug seeking in rats. J Neurosci 28: 8735–8739.

13. Liu Y, Postupna N, Falkenberg J, Anderson ME (2008) High frequency deep brain stimulation: what are the therapeutic mechanisms? Neurosci Biobehav Rev 32: 343–351.

14. Knapp CM, Tozier L, Pak A, Ciraulo DA, Kornetsky C (2009) Deep brain stimulation of the nucleus accumbens reduces ethanol consumption in rats. Pharmacol Biochem Behav 92: 474–479.

15. Doucette WT, Khokhar JY, Green AI (2015) Nucleus accumbens deep brain stimulation in a rat model of binge eating. Transl Psychiatry 5: e695.

16. Morrell MJ, Group RNSSiES (2011) Responsive cortical stimulation for the treatment of medically intractable partial epilepsy. Neurology 77: 1295–1304.

17. Rosin B, Slovik M, Mitelman R, Rivlin-Etzion M, Haber SN, et al. (2011) Closed-loop deep brain stimulation is superior in ameliorating parkinsonism. Neuron 72: 370–384.

18. Wu H, Miller KJ, Blumenfeld Z, Williams NR, Ravikumar VK, et al. (2017) Closing the loop on impulsivity via nucleus accumbens delta-band activity in mice and man. Proc Natl Acad Sci U S A.

19. Little S, Pogosyan A, Neal S, Zavala B, Zrinzo L, et al. (2013) Adaptive deep brain stimulation in advanced Parkinson disease. Ann Neurol 74: 449–457.

20. Carmichael JE, Gmaz JM, van der Meer MAA (2017) Gamma Oscillations in the Rat Ventral Striatum Originate in the Piriform Cortex. J Neurosci 37: 7962–7974.

21. van der Meer MA, Redish AD (2009) Low and High Gamma Oscillations in Rat Ventral Striatum have Distinct Relationships to Behavior, Reward, and Spiking Activity on a Learned Spatial Decision Task. Front Integr Neurosci 3: 9.

22. Tort AB, Komorowski RW, Manns JR, Kopell NJ, Eichenbaum H (2009) Theta-gamma coupling increases during the learning of item-context associations. Proc Natl Acad Sci U S A 106: 20942–20947.

23. Howe MW, Atallah HE, McCool A, Gibson DJ, Graybiel AM (2011) Habit learning is associated with major shifts in frequencies of oscillatory activity and synchronized spike firing in striatum. Proc Natl Acad Sci U S A 108: 16801–16806.

24. Doucette WT, Dwiel L, Boyce JE, Simon AS, Khokhar JY, et al. (2018) Machine learning based classification of deep brain stimulation outcomes in a rat model of binge eating using ventral striatal oscillations Front Psychiatry.

25. Berner LA, Avena NM, Hoebel BG (2008) Bingeing, self-restriction, and increased body weight in rats with limited access to a sweet-fat diet. Obesity (Silver Spring) 16: 1998– 2002.

26. Corwin RL (2004) Binge-type eating induced by limited access in rats does not require energy restriction on the previous day. Appetite 42: 139–142.

27. Corwin RL, Buda-Levin A (2004) Behavioral models of binge-type eating. Physiol Behav 82: 123–130.

28. Friedman J, Hastie T, Tibshirani R (2010) Regularization Paths for Generalized Linear Models via Coordinate Descent. J Stat Softw 33: 1–22.

29. Boggiano MM, Artiga AI, Pritchett CE, Chandler-Laney PC, Smith ML, et al. (2007) High intake of palatable food predicts binge-eating independent of susceptibility to obesity: an animal model of lean vs obese binge-eating and obesity with and without binge-eating. Int J Obes (Lond) 31: 1357–1367.

30. Castro DC, Cole SL, Berridge KC (2015) Lateral hypothalamus, nucleus accumbens, and ventral pallidum roles in eating and hunger: interactions between homeostatic and reward circuitry. Front Syst Neurosci 9: 90.

31. Urstadt KR, Stanley BG (2015) Direct hypothalamic and indirect trans-pallidal, trans-thalamic, or trans-septal control of accumbens signaling and their roles in food intake. Front Syst Neurosci 9: 8.

32. Shin AC, Pistell PJ, Phifer CB, Berthoud HR (2010) Reversible suppression of food reward behavior by chronic mu-opioid receptor antagonism in the nucleus accumbens. Neuroscience 170: 580–588.

33. Pecina S, Berridge KC (2000) Opioid site in nucleus accumbens shell mediates eating and hedonic ‘liking’ for food: map based on microinjection Fos plumes. Brain Res 863: 71–86.

34. de Araujo IE, Oliveira-Maia AJ, Sotnikova TD, Gainetdinov RR, Caron MG, et al. (2008) Food reward in the absence of taste receptor signaling. Neuron 57: 930–941.

35. Horn A, Reich M, Vorwerk J, Li N, Wenzel G, et al. (2017) Connectivity Predicts deep brain stimulation outcome in Parkinson disease. Ann Neurol 82: 67–78.

36. Shamir RR, Dolber T, Noecker AM, Walter BL, McIntyre CC (2015) Machine Learning Approach to Optimizing Combined Stimulation and Medication Therapies for Parkinson’s Disease. Brain Stimul 8: 1025–1032.

37. Heldman DA, Pulliam CL, Urrea Mendoza E, Gartner M, Giuffrida JP, et al. (2016) Computer-Guided Deep Brain Stimulation Programming for Parkinson’s Disease. Neuromodulation 19: 127–132.

38. Grahn PJ, Mallory GW, Khurram OU, Berry BM, Hachmann JT, et al. (2014) A neurochemical closed-loop controller for deep brain stimulation: toward individualized smart neuromodulation therapies. Front Neurosci 8: 169.

39. Kumar SS, Wulfing J, Okujeni S, Boedecker J, Riedmiller M, et al. (2016) Autonomous Optimization of Targeted Stimulation of Neuronal Networks. PLoS Comput Biol 12: e1005054.

40. Churchland MM, Yu BM, Cunningham JP, Sugrue LP, Cohen MR, et al. (2010) Stimulus onset quenches neural variability: a widespread cortical phenomenon. Nat Neurosci 13: 369–378.

41. Schurger A, Pereira F, Treisman A, Cohen JD (2010) Reproducibility distinguishes conscious from nonconscious neural representations. Science 327: 97–99.

42. Schurger A, Sarigiannidis I, Naccache L, Sitt JD, Dehaene S (2015) Cortical activity is more stable when sensory stimuli are consciously perceived. Proc Natl Acad Sci U S A 112: E2083–2092.

43. Ledberg A, Montagnini A, Coppola R, Bressler SL (2012) Reduced variability of ongoing and evoked cortical activity leads to improved behavioral performance. PLoS One 7: e43166.

44. Arazi A, Censor N, Dinstein I (2017) Neural Variability Quenching Predicts Individual Perceptual Abilities. J Neurosci 37: 97–109.

45. Saberi-Moghadam S, Ferrari-Toniolo S, Ferraina S, Caminiti R, Battaglia-Mayer A (2016) Modulation of Neural Variability in Premotor, Motor, and Posterior Parietal Cortex during Change of Motor Intention. J Neurosci 36: 4614–4623.

46. Churchland MM, Yu BM, Ryu SI, Santhanam G, Shenoy KV (2006) Neural variability in premotor cortex provides a signature of motor preparation. J Neurosci 26: 3697–3712.

